# An update to the database for *Acinetobacter baumannii* capsular polysaccharide locus typing extends the extensive and diverse repertoire of genes found at and outside the K locus

**DOI:** 10.1101/2022.05.19.492579

**Authors:** Sarah M. Cahill, Ruth M. Hall, Johanna J. Kenyon

**Affiliations:** Centre for Immunology and Infection Control, School of Biomedical Sciences, Faculty of Health, Queensland University of Technology, Brisbane, Australia; School of Life and Environmental Sciences, The University of Sydney, Sydney, Australia

**Keywords:** *Acinetobacter baumannii*, Kaptive, capsular polysaccharide, K locus, KL

## Abstract

2.

Several novel non-antibiotic therapeutics for the critical priority bacterial pathogen, *Acinetobacter baumannii*, rely on specificity to the cell-surface capsular polysaccharide (CPS). Hence, prediction of CPS type deduced from genes in whole genome sequence data underpins the development and application of these therapies. In this study, we provide a comprehensive update to the *A. baumannii* K locus reference sequence database for CPS typing (available in *Kaptive v. 2*.*0*.*1*) to include 145 new KL, providing a total of 237 KL reference sequences. The database was also reconfigured for compatibility with the updated *Kaptive v. 2*.*0*.*0* code that enables prediction of ‘K type’ from special logic parameters defined by detected combinations of KL and additional genes outside the K locus. Validation of the database against 8994 publicly available *A. baumannii* genome assemblies from NCBI databases identified the specific KL in 73.45% of genomes with perfect, very high or high confidence. Poor sequence quality or the presence of insertion sequences were the main reasons for lower confidence levels. Overall, 17 KL were overrepresented in available genomes, with KL2 the most common followed by the related KL3 and KL22. Substantial variation in gene content of the central portion of the K locus, that usually includes genes specific to the CPS type, included 34 distinct groups of genes for synthesis of various complex sugars and >400 genes for forming linkages between sugars or adding non-sugar substituents. A repertoire of 681 gene types were found across the 237 KL, with 88.4% found in <5% of KL.

**Significance as a BioResource to the community:** New therapies that target the bacterial polysaccharide capsule (CPS) show promise as effective tools to curb the high mortality rates associated with extensively resistant *A. baumannii*; one of the world’s most troublesome Gram-negative pathogens. As important information about the CPS structure produced by an isolate can be extracted from Whole Genome Sequences (WGS), simple bioinformatic tools and definitive sequence databases are needed to facilitate robust prediction of CPS type from WGS data. Here, we provide a comprehensive update to the international CPS sequence typing database for *A. baumannii*, increasing the utility of this resource for prediction of CPS type from WGS to assist with clinical surveillance, and/or the design and application of CPS-targeted therapies. This study is expected to further inform epidemiological tracking efforts, as well as the design of therapeutics targeting the CPS, enhancing global efforts to identify, trace and treat infections caused by this pathogen.

**Data summary:**

1. The updated *A. baumannii* KL reference sequence database including 241 fully annotated gene clusters is available for download under *Kaptive v. 2*.*0*.*1* at https://github.com/katholt/Kaptive.
2. Genome assemblies, short read data, or GenBank records used as representative reference sequence for each K locus are listed in Supplementary Table S1, and are referenced within each entry in the *A. baumannii* KL reference sequence database.

**The authors confirm all supporting data, code and protocols have been provided within the article or through supplementary data files**.

## 5. Introduction

Failure of antibiotic therapy due to the emergence of pan-resistant bacteria is a growing global health crisis. *Acinetobacter baumannii* is ranked as one of six leading bacterial species responsible for nearly three quarters of deaths associated with antibiotic resistance worldwide, with an estimated 80% of circulating isolates in many low- and middle-income countries resistant to last-line carbapenems [1]. Hence, new therapeutic options are urgently needed for treatment of carbapenem-resistant *A. baumannii*. Promising strategies include monoclonal antibodies or bacteriophage [2]. Both strategies involve binding to cells via interaction with exposed structures on the bacterial cell surface, and can display specificity for structural epitopes of the polysaccharide capsule (CPS). However, in this species, even closely related isolates can produce different forms of CPS, making knowledge of the specific CPS type in the infection to be treated critical. Hence, the ability to determine CPS type is needed to underpin the design and application of these therapies. The genetics underlying the CPS type has also proven valuable as an epidemiological marker [3-7]. Finally, recent studies have associated some specific CPS types [8, 9] or alterations in the CPS structure [10] with increased virulence. Hence, the determination of the specific type produced in problem strains is important in several areas.

As information about CPS type can be deduced from genes in bacterial genomes, whole genome sequencing (WGS) is an attractive approach for CPS typing that is more readily accessible than traditional laboratory-based serological typing methods. For *A. baumannii*, most of the genes responsible for CPS biosynthesis are clustered together in the chromosome between *fkpA* and *lldP* genes [11]. However, many different sets of genes have been found at this ‘K locus’ (KL). To facilitate their identification, 92 fully annotated KL reference sequences were recently compiled into a curated database and released publicly [12]. The database is compatible with the bioinformatics search tool, *Kaptive* [13] and *Kaptive*-Web [14]. This database was validated against 3415 genome sequences available in the NCBI non-redundant and WGS databases at that time and 642 genomes assembled from reads available in NCBI SRA database. However, it was noted that additional KL configurations were known, and that there may be many more KL yet to be documented [12]. In fact, more than 128 distinct K loci were known at the time and an additional 78 KL have since been identified as additional sequence data became available [15, 16, 17 and Kenyon, unpublished data].

All known gene clusters at the *A. baumannii* K locus follow a general pattern that includes 3 ‘regions’ [11, 12]. Region 1 always consists of essential CPS export genes (*wza, wzb* and *wzc*) that are transcribed in the opposite direction to the remainder of the locus.Region 2, the central portion, includes many different sets of genes and these determine the composition and structure of the K unit making up the specific CPS type. Region 3 flanks the other side of Region 2, and always includes genes for the synthesis of simple sugar precursors (*galU, ugd, gpi, pgm*), though genes can be variably inserted between *gpi* and *pgm*. The *gne1* gene for D-Gal*p* or D-Gal*p*NAc synthesis is often present between *gpi* and *pgm* [11] but has been found to be absent from some KL that do not include D-Gal*p* or D-Gal*p*NAc in the corresponding CPS [17-22]. Other genes can also be found in this position giving rise to some variation in Region 3 [19, 23-26].

In general, a unique KL identifier is assigned to a sequence when there is a detectable difference in gene content with genes identified based on a product sequence identity cut-off of 85%. However, as chemical structures of CPS produced by 70 distinct KL have been determined [e.g. 17, 19, 20, 27-34], it is now known that differences in the ‘conserved’ genes in Region 3 (i.e. *galU, ugd, gpi, pgm*) do not influence the type of CPS produced. In addition, variation in the genes in Region 1 (*wza, wzb, wzc)* does not affect their essential role in capsule export. Therefore, a new KL number is only assigned to a sequence when there is a detectable difference in genes in Region 2 and/or the variable portion of Region 3 (between *gpi* and *pgm*).

In most cases, complete correlation between the genetic content of specific KL and the structural features of the corresponding CPS type have been reported. However, for a few strains, the *wzy* gene has been shown to be missing from Region 2 of the K locus or interrupted by an insertion sequence (IS), and a replacement *wzy* gene was found to the left of Region 1 as in KL8 [11] or in a defined genomic island outside of the K locus [15, 35, 36]. In a recent study, an additional Wzy polymerase gene was identified in prophage sequence integrated elsewhere in the chromosome, and was found to alter the linkage between oligosaccharide K-units that make up the CPS structure [37]. Acetyltransferase genes with encoded products that have been shown to modify the CPS by acetylation have also been found in integrated phage genomes [38]. Therefore, detection of these additional genetic determinants in the genome will be essential to achieve robust prediction of CPS type from WGS data.

Recently, the *Kaptive* code was updated (*Kaptive v. 2*.*0*.*0*) to include an additional function that was designed specifically for discrimination of O-antigen serotypes in the *Klebsiella pneumoniae* species complex [39]. For this function, determination of serotype or ‘type’ is based on the detection of either the O-antigen locus (OL) type alone or a defined combination of OL and ‘extra genes’ in a genome assembly known to be involved in the determination of a specific serotype. As an active CPS serotyping scheme does not exist for *A. baumannii*, ‘type’ has previously been used to refer to the chemical structure of the CPS produced by an isolate as defined by the KL number i.e. the KL2 sequence produces the K2 type CPS [32]. In cases where genes outside the K locus have been shown to modify the CPS type, a suffix is now added to the K type name. For example, this was recently done for K127-Wzy_Ph1_, which is defined as the structure formed by KL127 and Wzy encoded in prophage (Ph) [37]. Other examples, such as K19 and K24 modified by Wzy proteins encoded by genomic islands, GI-1 and GI-2, respectively, have also now been renamed to indicate the role of extra-KL genes. Hence, the new *Kaptive v. 2*.*0*.*0* function can be harnessed to predict *A. baumannii* CPS ‘type’ as defined by structural data where this is available.

In this study, we provide a comprehensive update to the *A. baumannii* CPS reference sequence database to include the known KL not included in the original version and new KL sequences detected in 8994 publicly available *A. baumannii* genome sequences. Special logic parameters to enable prediction of the CPS type based on KL or the detected combination of a specific KL with ‘extra genes’ have also been included as well as information relating to K type where structures have been determined. The updated database was validated against the same large genome set and a smaller set of complete genomes. A detailed assessment of gene repertoire at the chromosomal K locus was also conducted.

## 6. Methods

### *A. baumannii* genome assemblies

A total of 9065 genome assemblies listed under the *Acinetobacter baumannii* taxonomic classification in the NCBI non-redundant and WGS databases (10^th^ June, 2021) were downloaded for local analysis. Assemblies were first assessed for the presence of the *A. baumannii-*specific *oxaAb* gene (also known as *bla*_OXA-51-like_; available in *A. baumannii* strain A1 complete genome sequence under GenBank accession number CP010781.1, base positions 1753305 to 1754129) with BLASTn using a cut-off of >90% combined coverage with >95% nucleotide sequence identity to confirm the *A. baumannii* species assignment (as previously defined in [12]). Only confirmed *oxaAb-*positive genomes were used for downstream analyses.

### Identification and annotation of novel K locus sequences

*A. baumannii* genome assemblies (n = 8994) were screened against the original version of the *A. baumannii* KL reference database [12] available in the *Kaptive versions 0*.*7*.*0-2*.*0*.*0* (https://github.com/katholt/Kaptive) and then using an extended in-house database of 206 KL (*unpublished*) with the command-line version of *Kaptive v 0*.*7*.*0* [13]. The search was conducted using a parameter defining the “minimum gene identity” cut-off as 85% as is standard for *A. baumannii* KL typing [11]. Output results from the in-house screen for matches with a reported confidence level less than ‘perfect’ were examined. Matches with length discrepancies, additional or missing genes, or those with <95% coverage and/or <95% nucleotide sequence identity to the best matched reference sequence were manually inspected to identify novel gene clusters.

Novel KL were annotated using the established nomenclature system for *A. baumannii* K locus typing [11,12]. Briefly, KL were assigned a new number if any differences were detected in gene presence/absence in Region 2 and between *gpi* and *pgm* genes in Region 3, and standard gene names were used to indicate enzyme function. For glycosyltransferase (*gtr*), acetyl or acyltransferase (*atr*), and pyruvyltransferase (*ptr*) genes where sequence differences may result in a change of substrate preference, new numbers were assigned when the product of the gene had <85% amino acid (aa) sequence identity to the closest match.

KL comparisons were generated using EasyFig *v 2*.*2*.*2* [40], and genome comparisons assembled using Mauve *v 2*.*4*.*0* [41] to order contigs, followed by BRIG [42] to generate a circular comparison. Where necessary, read quality was assessed using FASTQC *v 0*.*11*.*9* (https://www.bioinformatics.babraham.ac.uk/projects/fastqc/) and assembly quality examined using QUAST *v. 5*.*02* [43]. For some cases, multi locus sequence typing (MLST) was performed on the assembly using both Oxford and Institute Pasteur schemes established for *A. baumannii* in PubMLST with the MLST package (https://github.com/tseemann/mlst).

### Curation of updated *A. baumannii* KL reference sequence database

A GenBank format file (.gbk) for each new locus sequence was prepared, which included the nucleotide sequence and annotations of all coding sequences in the locus. Where the only available representative of a KL included an insertion sequence (IS), we substituted the sequence with a manually generated version where the IS and target site duplication were removed in order to include a KL that represents the presumptive ancestral, non-modified sequence as is required for accurate typing by *Kaptive*. This was the case for 23 different KL, which are indicated in Supplementary Table S1. An additional note field was added for all reference loci to define the K type where structural data for the CPS was available for the specific KL sequence as indicated in the reference record in each entry in the database. In cases where no structural data is available, the note field specifies the type as unknown.

For CPS structures known to be modified by additional genes outside the K locus, the note field indicates ‘special logic’ to be applied by *Kaptive*. This directs the tool to perform an additional tBLASTn search for ‘extra genes’ supplied in the database, and an additional “Acinetobacter_baumannii_k_locus_primary_reference.logic” file then specifies ‘type’ when a specific combination of KL and extra genes is found. GenBank records for six ‘extra genes’ added to the database include: *wzy* and *atr25* genes in ‘genomic island 1’ (GI-1) involved in K19 synthesis [35]; *wzy* in ‘genomic island 2’ (GI-2) for K24 synthesis [36]; *atr29* and *atr30* in prophage (Ph) found to modify the K46 and K5 structures, respectively [38]; and *wzy* in prophage that has recently been found to modify the K127 CPS [37].

All GenBank-format .gbk files were concatenated into a multi-record file to produce an updated KL reference database for release as *Kaptive v. 2*.*0*.*1*. The updated database was integrated with the *Kaptive-Web* platform (http://kaptive.holtlab.net/), and is available for download from https://github.com/katholt/Kaptive for use with the command-line *Kaptive v. 2*.*0*.*0*. The database was validated on the same genome pool using *Kaptive v. 2*.*0*.*0* with a parameter defining the minimum gene product identity cut-off as 85%.

### Analysis of sequence features and gene frequency

To generate an overview of the sequence lengths and total repertoire of genes found at the *A. baumannii* K locus, Prokka *v. 1*.*13* [44] was used to generate gff3 files for each individual K locus sequence using manual annotations available from each gbk record. The complete length of each gene cluster, along with the total number of open reading frames, and number of *gtr* genes was manually tabulated. The gff3 files were then used as an input for the pan genome tool Roary *v. 3*.*13*.*0* [45] to generate a gene presence/absence matrix using a cut-off parameter of 85% aa sequence identity. The matrix was used to determined homology groups defined as genes encoding products with >85% aa sequence identity and >330 bp in length. Summarised data was visualised using the ggplot2 package in RStudio *v. 1*.*2*.*5033* [46].

## 7. Results

**Screening for novel CPS biosynthesis gene clusters at the K locus**

A total of 9,065 genome assemblies available under the *Acinetobacter baumannii* taxonomy classification (Taxonomy ID: 470) were downloaded from NCBI GenBank and WGS databases. The intrinsic *oxaAb* gene could not be identified in 71 assemblies, hence these were excluded from further analysis. Confirmed *A. baumannii* genome assemblies (n=8994) were screened against the original *A. baumannii* KL reference sequence database [12] included in *Kaptive* versions *(v) 0*.*7*.*0-2*.*0*.*0* (hereafter referred to as database *v 0*.*7*.*0-2*.*0*.*0*). The confidence levels (categorical measure of match quality) called by *Kaptive v 0*.*7*.*0* using this database were: 794 (perfect), 4794 (very high), 443 (high), 1776 (good), 192 (low) and 995 (none) (Supplementary Table S2; summarised in Table 1). This revealed that 62.13% of genome assemblies could be confidently assigned a match indicated by a confidence level of ‘Perfect’ (the identified locus is in a single contiguous sequence that shares 100% coverage and 100% nucleotide sequence identity) or ‘Very high’ (a single contiguous sequence sharing ≥99% coverage and ≥95% nucleotide sequence identity with the best match reference sequence with no additional and/or missing coding sequences).

**Table 1.** Summary of *Kaptive* search results.

|  | Match Confidence Scores |  |  |  |  |  |  |
| --- | --- | --- | --- | --- | --- | --- | --- |
|  | Perfect | Very high | High | Good | Low | None | Total |
| Database v 0.7.0-2.0.0 | 794 | 4794 | 443 | 1776 | 192 | 995 | 8994 |
| Database; in-house | 922 | 5124 | 449 | 1726 | 135 | 638 | 8994 |
| Database v 2.0.1 | 1012 | 5158 | 436 | 1647 | 97 | 644 | 8994 |

Since additional KL reference sequences have been characterised following the release of the first database in 2020, the genome pool was reanalysed using an extended in-house database that includes a total of 206 KL made up of the 92 KL reference sequences in the original database plus the 36 previously characterised KL that were not included in the original database, and an additional 78 distinct KL characterised since this time [15, 16, 17 and Kenyon, unpublished data]. Confidence levels obtained were: 922 (perfect), 5124 (very high), 449 (high), 1726 (good), 135 (low) and 638 (none) (Table 1; Supplementary Table S3). This secondary screen revealed a shift in the number of assemblies in each confidence level with the proportion of matches scored ‘perfect’ or ‘very high’ confidence rising slightly to 67.22%. Given that a match with ‘perfect’ confidence indicates identity to the reference sequence, only matches assigned with a confidence level of ‘very high’ or less were further examined to identify novel locus sequences.

### ‘Very high’ confidence matches are close relatives or IS variants of reference sequences

Manual inspection of the output data from the screen using the in-house database of 206 KL revealed that 5124 assemblies had a match assigned with ‘very high’ confidence. Of these 5124 assignments, 5094 (99.4%) were considered very close relatives of the best match locus with single nucleotide polymorphisms (SNPs). However, 30 (0.6%) had a discrepancy in the total length of the locus match with >700 bp of additional sequence. For 28 of these 30 assignments, the additional sequence was found to be an insertion sequence (IS), and thus these loci were deemed variants of the archetypal KL reference sequence in the database. The remaining two KL with discrepant lengths had variable-length insertions of ‘N’ bases, which indicated possible issues with sequence or assembly quality.

### ‘High’ confidence matches include IS variants and novel KL sequences

A total of 449 assemblies were assigned a match with a confidence level of ‘high’ indicating that the KL was found in a single piece with ≥99% coverage but less than three missing gene products and no extra genes. For 434 of 449 assignments (96.66%), the locus sequence had been correctly identified but with detectable problems. For 415 of these, the locus match had ≥99% coverage, ≥89% nucleotide sequence identity and no more than +/- 101 bp of discrepancy in sequence length to a KL reference sequence in the database, though SNPs or base insertions/deletions resulting in frameshifts in known coding sequences were found. For the other 19 assignments, significant discrepancies in the total sequence length of the match (greater than +/-101 bp) were reported by *Kaptive*. For 15 of these, the additional sequence was confirmed to be one or more IS insertions. Three others had strings of missing bases, whereas one had a string of additional ‘N’ bases. Hence, 19 were considered variants or possible variants of the best match KL reference sequence. Therefore, the specific KL had been correctly identified in 72.05% of genomes assigned with either ‘perfect’, ‘very high’ and ‘high’ matches.

For 11 of the remaining 15 ‘high’ assemblies, nucleotide sequence identity to a best match locus of KL33 was <94%. However, the expected *psaD* and *psaE* genes that encode a nucleotidase and an acetyltransferase involved in the synthesis of 5,7-di-*N*-acetylpseudaminic acid, respectively, were reported missing by *Kaptive*. Analysis of these genome assemblies revealed that they all carried the same sequence at the K locus, which differed from KL33 only in a small segment where the *psaD* and *psaE* genes of KL33 are replaced by two related but novel genes, designated here as *psaI* and *psaJ* (Fig. 1A). Both the encoded PsaI and PsaJ products share 78.9% aa sequence identity to their PsaD and PsaE homologues. Therefore, the KL sequence was considered novel and designated KL235. The *psaI* and *psaJ* genes were also identified in place of *psaD/psaE* in a further 2 of the 15 assemblies, which both had >95% identity to KL121. These were also considered novel and designated KL218 (Fig. 1A). Though PsaI and PsaJ may produce the same sugar product as PsaD and PsaE, it is possible that the difference in sequence could result in a new acylated derivative of pseudaminic acid, and structural studies will be needed to confirm this.

**Figure 1.**
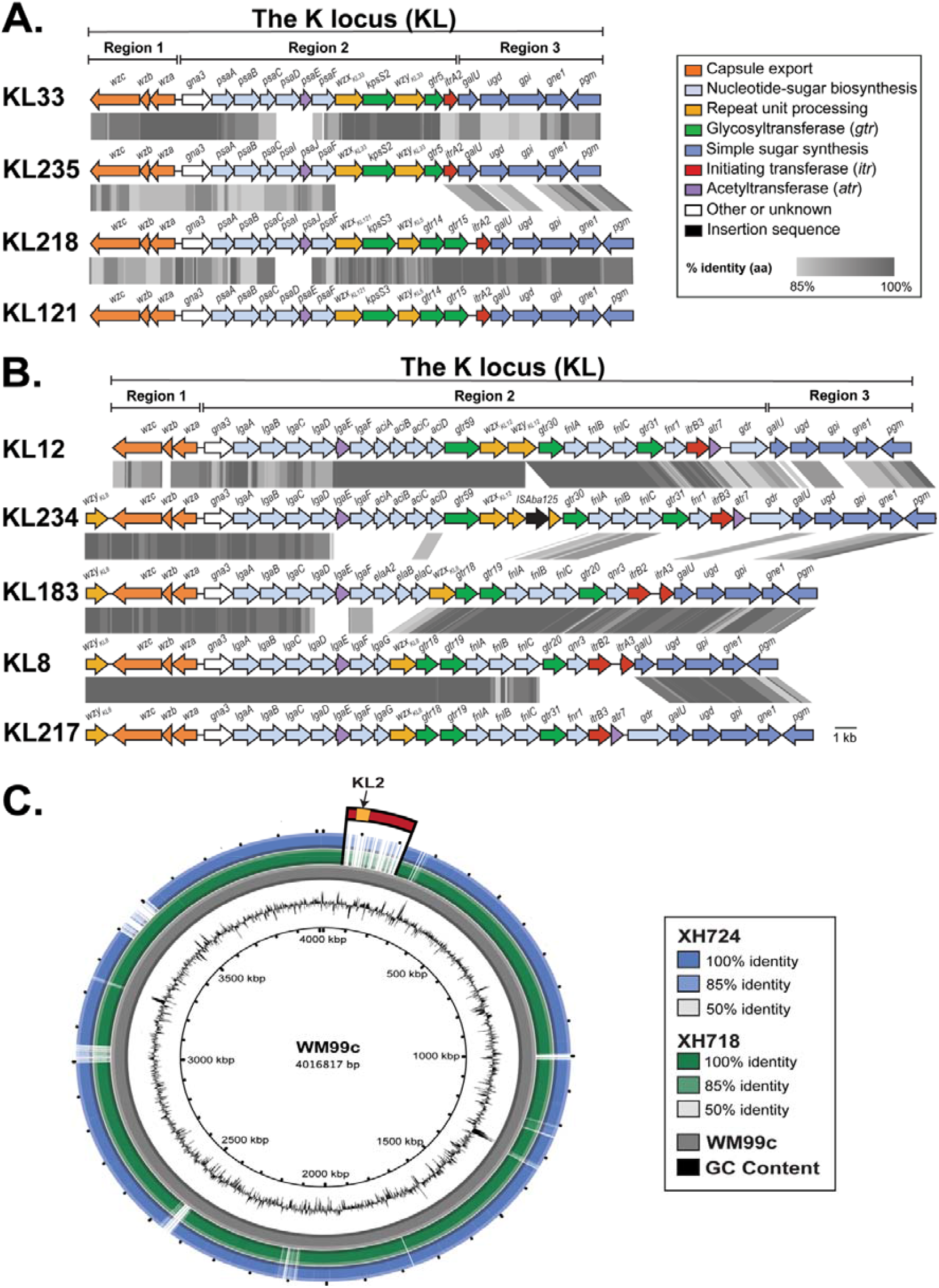
**(A)** Comparison of KL33 to the best match reference sequence, KL235, and related sequences, KL218 and KL121. **(B)** Comparison of KL234 to the best match reference sequence, KL12, and to related sequences with *wzy* downstream of *wzc*. Genes are coloured to the functional category of their gene products with legend shown top right. Grey shading between gene clusters indicates amino acid sequence identities with scale shown in the legend. Figures drawn to scale using Easyfig [40] and annotated/coloured in Adobe Illustrator. **(C)** BRIG multiple sequence alignment of genomes from strains XH724 and XH718 aligned to the WM99c reference genome (NCBI accession number CP032055.1). Contigs were reordered using MAUVE [41] prior to BRIG [42]. Location of the KL2 locus in the WM99c reference genome is marked orange.

The final two assemblies (NCBI assembly accession numbers GCA_005280695.1 and GCA_013305465.1) had been assigned a best match locus of KL12 with 100% sequence coverage and 94.7% nucleotide sequence identity by *Kaptive*. However, the output indicated an additional 1083 bp of sequence, and the expected gene coding for the Wzy polymerase was missing. Manual inspection of the two genome assemblies revealed that they carried the same K locus sequence, and direct comparison of these DNA sequences to KL12 (Fig. 1B) revealed that the *wzy*_*KL12*_ gene was present but interrupted by an IS*Aba*125 insertion sequence. One of these assemblies, GCA_013305465.1, had been reported in a clinical isolate from Australia [47], and an additional gene sharing 100% identity with *wzy* from KL183 in the database was identified in the *Kaptive* output field, ‘Other genes outside locus’. However, this *wzy* gene is in fact in the locus (i.e. between *fkpA* and *lldP*) but in an unusual position between *fkpA* and *wzc* at the 5’-end of the locus (Fig. 1B). The location of *wzy* at the beginning of the locus adjacent to *fkpA* was previously reported for KL8 [11], and the *wzy* gene from KL183 is identical to the KL8 *wzy*. A similar configuration was also found for KL217 (see below). Hence, this gene was assigned the name *wzy*_*KL8*,_ and KL234 was assigned to this novel locus.

### ‘Good’, ‘Low’ or ‘None’ confidence matches

A total of 2499 genome assemblies were assigned a match with a ‘good’ (1726), ‘low’ (135) or ‘none’ (638) confidence level. These included only 231 (9.24%) with a best match locus found in a single contiguous sequence, and 2268 genomes (90.75%) found in two or more pieces (indicated by a ‘?’ problem score in the *Kaptive* output). As detectable breaks in KL loci often suggest that a genome sequence or an assembly is poor quality or that loci are variants in which an IS has interrupted the KL sequence, the 2268 assemblies with loci found in more than 1 piece were not further investigated. For the 231 contiguous sequences, 135 loci (76 ‘good’, 24 ‘low’ and 35 ‘none’) were found to include numerous SNPs relative to the assigned reference sequence or insertions of ‘N’ bases suggesting problems with sequence or assembly quality. A further 46 ‘good’ matches included an IS indicating these were variants of the best match reference sequence.

Of the 50 remaining contiguous matches, four genome assemblies were found to be missing significant portions of the K locus sequence. One of these had a match confidence of ‘good’ and was missing 13% of the assigned KL124 locus sequence, while a second locus had a match confidence of ‘none’ and was missing 42% of the assigned KL13 locus. Another two genome assemblies (GCA_001862175.1 and GCA_001862305.1) were assigned a best match to KL92 with a ‘none’ confidence level and 0 of 22 expected genes identified. Manual inspection of the two genome assemblies and comparison to the complete genome sequence of a related strain revealed that both assemblies had a ∼150 kb deletion that included the K locus (Fig. 1C). The associated NCBI assembly data indicated these were clinical isolates sequenced using Illumina Hiseq 2000 and assembled using CLC Genomic workbench *v*.

*8*.*5*.*1*. To assess if the deletion may be due to poor assembly or read quality, genome assemblies were subjected to QUAST, and their short reads (SRR3381523 and SRR3381529) to FastQC. Results outputs suggested good read and assembly quality (<50 contigs, length= 3.6-3.7 Mbp, 38.9% GC), and these genome assemblies were not further investigated. These are surprising findings that arise from the fact that the current version of *Kaptive* still assigns a best match KL even when the sequence is not present, and this will need to be addressed in a future update to the *Kaptive* code.

The remaining 46 genome assemblies (29 ‘good’, 2 ‘low’, 15 ‘none’) were found to have <95% DNA sequence coverage, <95% DNA sequence identity, significant length discrepancies (>400 bp), missing expected genes and/or presence of unexpected genes in the locus sequence. Amongst the 46 assemblies, 28 novel KLs were identified by manual inspection. 27 of these KL were found to follow the same general pattern as for other gene clusters described at the *A. baumannii* K locus to date, in that they consisted of three defined regions with one *wzx* gene and one *wzy* gene in Region 2 of each gene cluster. The exception, KL217, included the *wzy*_*KL8*_ gene in the location at the start of the K locus as described for KL234 (see Fig. 1A). Therefore, together with KL218, KL234 and KL235 described above, a total of 31 novel KL were identified amongst the 8994 genomes studied, bringing the total number of known KL to 237.

### Annotation of novel genes

Annotations were manually curated in accordance to the standard nomenclature system for *A. baumannii* CPS biosynthesis genes [11,12] for 145 KL, which included the novel 31 KL detected above, as well as the 114 not included in the previous version of the database or identified since the first release. Several novel genes and gene modules were identified across the 145 types and are described in further detail below.

Amongst the 145 additional KLs to be included in the new iteration of the database, a total of 75 genes were predicted to encode novel glycosyltransferases (defined as <85% aa identity to known types) not seen in the previous database. The products of three of these were found to be homologues of glycosyltransferases previously annotated as KpsS1 and KpsS2, and hence the genes were named *kpsS3-kpsS5* consistent with the nomenclature used previously for this Gtr type [11, 32, 48]. All other predicted glycosyltransferases were assigned new *gtr* numbers. Similarly, 19 genes were predicted to encode new acetyl-/acyl-transferases and were assigned new *atr* names, while four new putative pyruvyltransferase genes were found and assigned *ptr* names. In addition, 12 genes of unknown function (*orf*) were also identified and further work will be needed to determine if these play a role in CPS biosynthesis.

Several novel genes likely to be involved in the synthesis of a monosaccharide were also found. For 6 KL (KL62, KL79, KL97, KL110, KL183, KL192), a homologue of the *elaA* gene, designated *elaA2*, was identified adjacent to *elaBC* and in place of *elaA*. ElaA is a putative oxidoreductase involved in the synthesis of 8-epilegionaminic acid (8eLeg) in the K49 CPS structure [31]. As ElaA and ElaA2 share 84% aa sequence identity, they likely catalyse the same reaction. However, structural studies of the CPS produced by these 6 loci will be needed to assess if ElaA2 also produces 8eLeg or a related sugar. Other additional homologues of sugar synthesis genes were identified for *mnaA* and *dmaA*, and these genes are predicted to be involved in the synthesis of UDP-D-Man*p*NAc and UDP-2,3-diacetamido-2,3-dideoxy-D-mannuronic acid, respectively. Gene homologues (encode proteins <85% identical) were assigned numbers (*mnaA1-mnaA4* and *dmaA1-dmaA4*) and further structural studies of the CPS will also be needed to confirm the type of sugar(s) produced.

Two genes located adjacent to each other in seven different KL (KL126, KL207, KL208, KL209, KL219, KL228, KL236; Fig. 2A) were found to encode homologues (>85% aa identity) of WeeE and WeeF from *Acinetobacter venetianus* RAG-1 ‘emulsan’ gene cluster (GenBank accession number AJ243431.1). WeeE and WeeF have previously been postulated to be involved in the synthesis of UDP-N-acyl-L-galactosaminuronic acid (UDP-L-Gal*p*NAcA), which is present in the RAG-1 CPS [49]. As in the nomenclature system for *A. baumannii* CPS biosynthesis gene clusters are designated names after the putative sugar product or function of the encoded enzyme in the synthesis pathway [11], the two genes were named *gnlA* and *gnlB* for UDP-L-Gal*p*NAcA, rather than *weeE* and *weeF*. The *gnlB* gene was found without *gnlA* in an eighth gene cluster, KL215.

**Figure 2.**
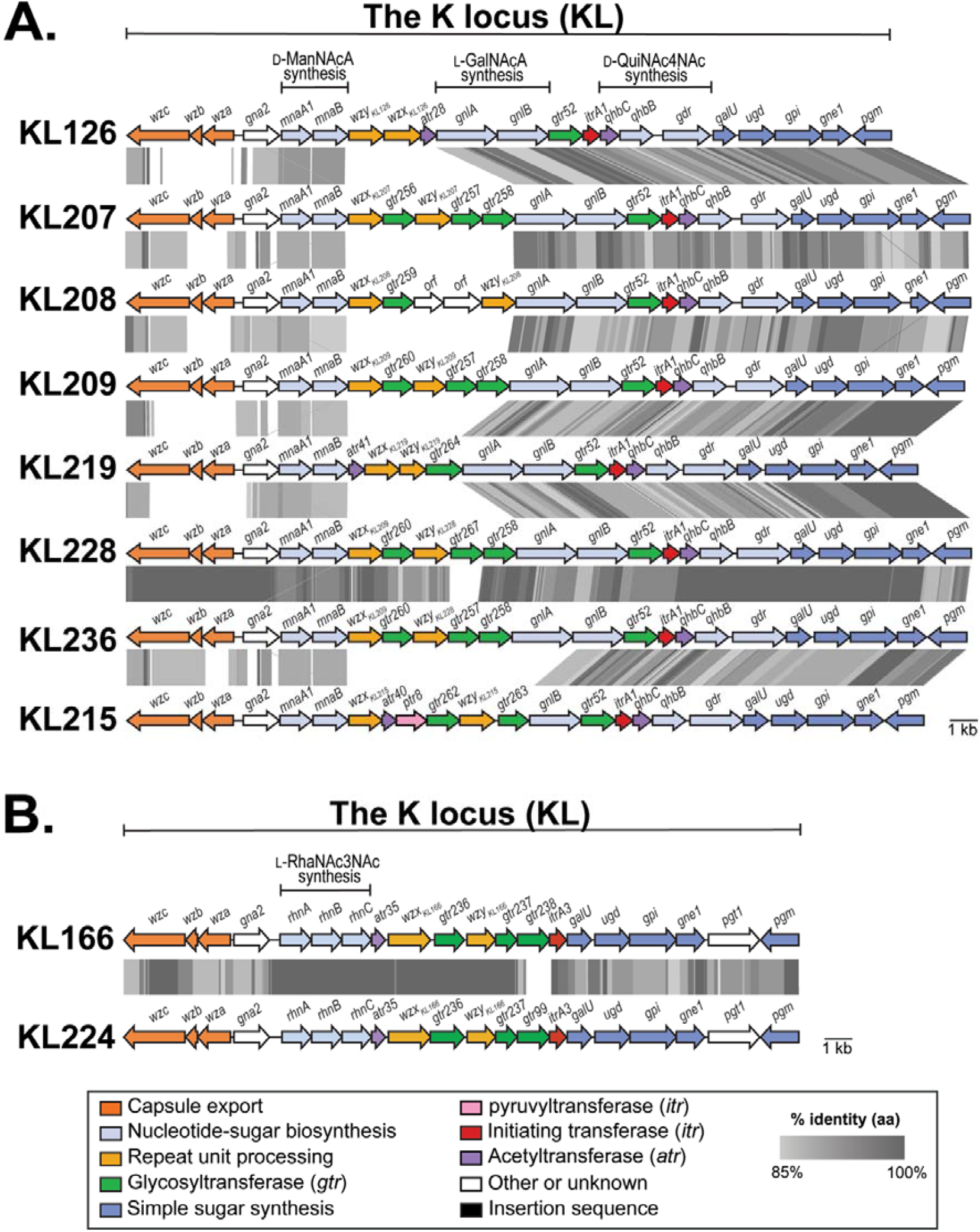
Comparison of *A. baumannii* KL that include **(A)** novel *gnlA* and/or *gnlB* genes, and **(B)** novel *rhn* genes. Genes are coloured to the functional category of their gene products with colour legend shown below. Grey shading between gene clusters indicates amino acid sequence identities with scale shown in the legend below. Figures drawn to scale using Easyfig [40] and annotated/coloured in Adobe Illustrator.

Three novel genes found in KL166 and KL224 (Fig. 2B) encode proteins sharing 33-66% aa identity with WeiS, WeiP, and WeiQ from the *Escherichia coli* O109 O-antigen gene cluster (GenBank accession number HM485572.1). WeiS, WeiP, and WeiQ have been predicted to be involved in the synthesis of 2,3-diacetamido-2,3,6-trideoxy-L-mannose (L-RhaNAc3NAc) found in the O109 structure [50]. As for *gnlAB*, the three genes were assigned new names, *rhnA, rhnB* and *rhnC* for L-<u>Rh</u>a<u>N</u>. Hence, the *gnl* and *rhn* names were added to the nomenclature scheme for *A. baumannii* CPS biosynthesis genes and an updated list of name descriptors can be found in Table 2.

**Table 2.**
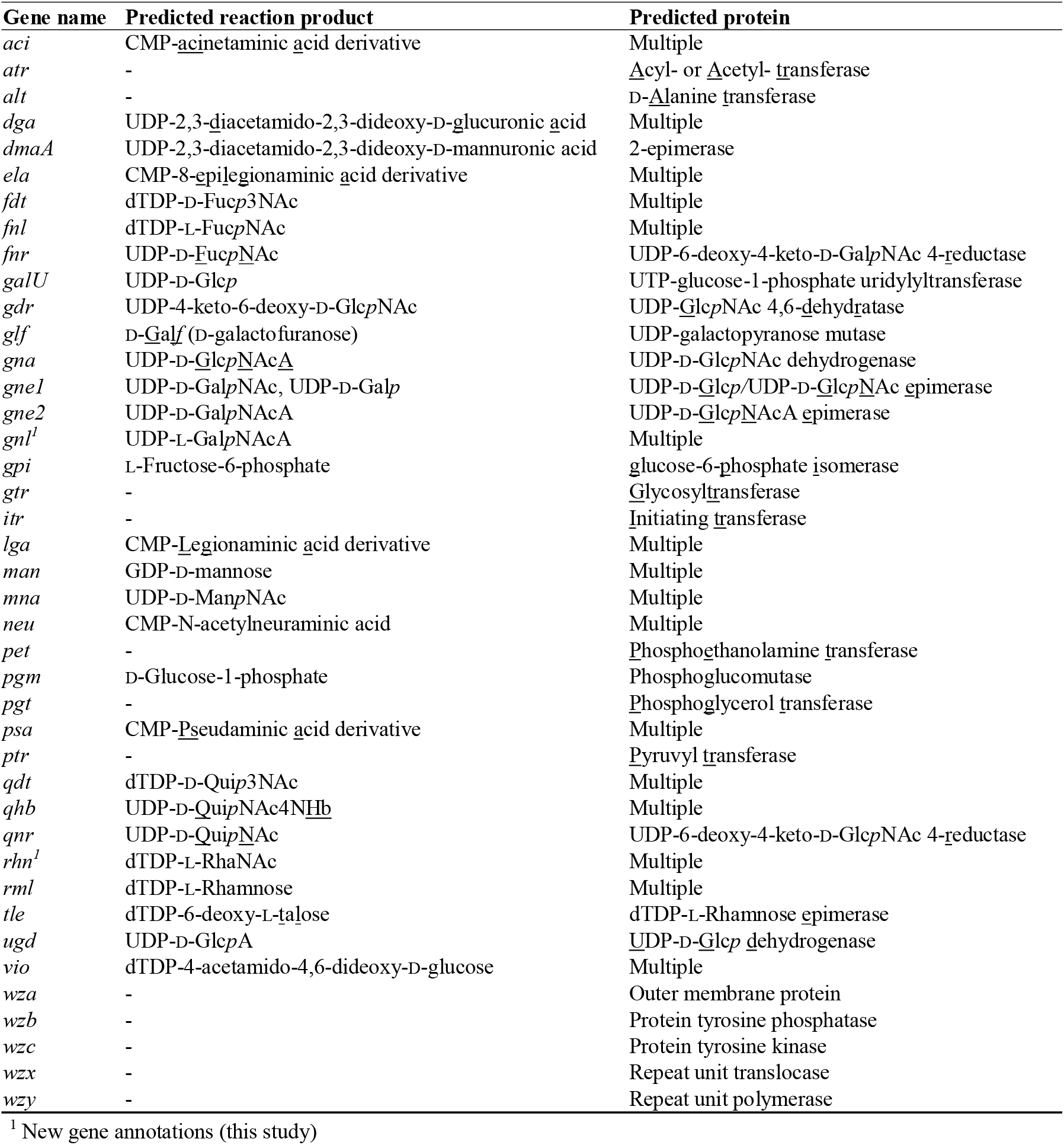
Updated gene nomenclature key for *A. baumannii* K loci.

### Updating the *A. baumannii* KL reference sequence database

A representative of each novel KL sequence was chosen for addition to the reference database. Fully annotated locus sequences were curated into GenBank format files (.gbk), then added to the multi-record database to create a new iteration that includes a total of 237 KL. Two reference sequences, KL38 and KL78, were retained in the updated database although the isolates carrying them have since been reassigned to other *Acinetobacter* species and are no longer considered *A. baumannii* isolates. The KL38 and KL78 representative reference sequences will be replaced with an *A. baumannii* sequence if one is later identified in the species. Information on the representative sequences chosen for the database are available in Supplementary Table S1.

As the latest release of the *Kaptive* code (*v. 2*.*0*.*0*) includes a new function that enables ‘type’ to be inferred from locus sequence, an additional note field was added for all reference loci to define the ‘type’ of CPS structure produced by a given KL. Citations for associated structural data were also integrated into the database in the reference section of each record, so that users are referred to the relevant publication(s). Where a structure for a specific KL has not yet been determined, the integrated note field defines the ‘type’ as unknown.

For CPS that have been found to require or be modified by additional genes located outside the K locus, the added note indicates that ‘special logic’ needs to be applied by *Kaptive*. This directs the tool to perform an additional tBLASTn search for ‘extra genes’ supplied in the database file. If detected, an additional “Acinetobacter_baumannii_k_locus_primary_reference.logic” file then specifies the ‘type’ when a specific combination of KL and extra genes are found. This feature will be further developed in future updates.

### Validation of the updated *A. baumannii* KL reference sequence database

To assess the utility of the updated KL reference sequence database (hereafter referred to as *v. 2*.*0*.*1*) with the constructed special logic file, the pool of 8994 genome assemblies were re-screened using *Kaptive v. 2*.*0*.*0* using the same parameters. The output confidence levels were: 1012 (perfect), 5158 (very high), 436 (high), 1647 (good), 97 (low) and 644 (none) (Table 1; Supplementary Table S4). Hence, the percentage of genome assemblies that could be typed with a confidence score of ‘perfect’, ‘very high’, or ‘high’ rose to 73.45% (Fig. 3A). As a greater number of KL were confidently assigned with the updated database, there was an observed decrease in the overall number of ‘problem’ scores called by *Kaptive*. This included a decrease of 627 assemblies with missing expected genes (denoted in the output by a ‘-’ symbol), 375 assemblies with additional genes (‘+’), and 418 assemblies with one or more expected genes with low identity (‘*’). An unexpected decrease was observed for matches found in more than one piece (‘?’), with 30 fewer assemblies reported with this problem score using *Kaptive 2*.*0*.*1* (Fig. 3B).

**Figure 3.**
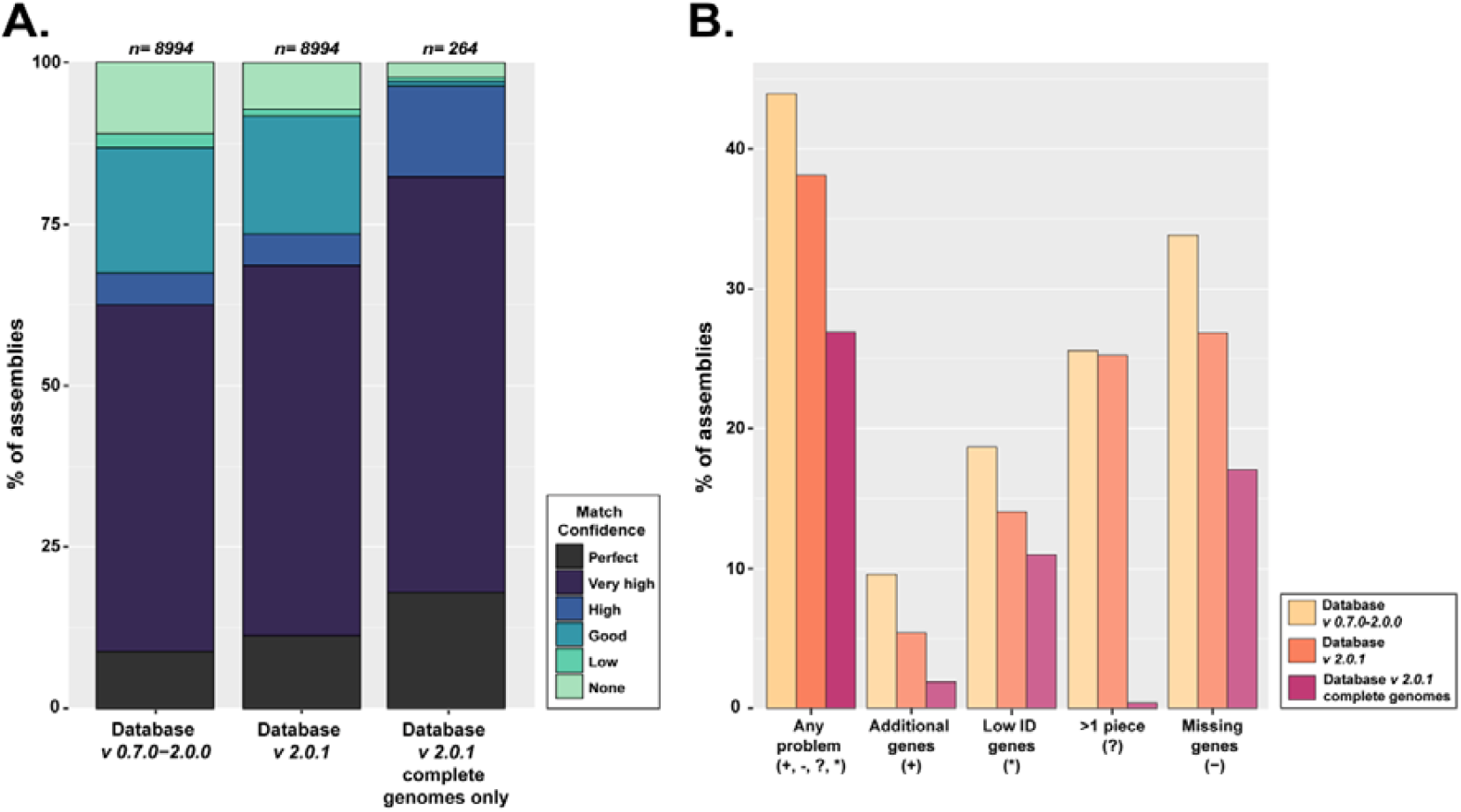
Comparison of *Kaptive* performance using the original and updated *A. baumannii* KL reference sequence databases. **(A)** Comparison of match confidence scores visualised as a stack bar blot. Database version is indicated below, and the number of genomes assessed per column is shown above. Colours indicate match confidence scores, and the key is shown on the right. Match confidence is a categorical measure of match quality between query and reference sequence in the database, and definitions for each category can be found in [12]. (**B)** Comparison of ‘problem’ scores visualised as a bar plot. Database version is indicated by the colour key shown on the right, and type of problem score is shown below. Figures were created using ggplot2 package in RStudio [46].

### Assessment of complete genome sequences

To further evaluate the impact of sequence quality on the number of problem scores called by *Kaptive*, results for 264 completed genome sequences with the chromosome available in a single contig were extracted for more detailed analyses. The match confidence scores, length discrepancies, and problem scores were summarised for the 264 complete genomes, and are shown in Table 3. More than 80% of the completed genomes were assigned a match confidence score less than ‘perfect’ (Fig. 3A). For these, all matches with a detected length discrepancy of >500 bp (20 total) were found to include an IS insertion. A total of 71 complete genomes were assigned at least one problem score (Fig. 3B). However, the majority were due to low detectable identity of some genes, frameshifts or better sequence matches of the same gene found in a different KL reference sequence in the database. Interestingly, one complete genome received a ‘?’ problem score suggesting that the K locus was not in a single piece. For this genome, the sequence had been opened within the K locus rather than at the origin of replication, leading to detection of the K locus at both the start and the end of the opened chromosome.

**Table 3.** Summary of *Kaptive v 2*.*0*.*1* results for complete genomes only.

| Confidence score | Total number of assemblies | Matches with length discrepancy (> +/- 500bp) | Matches including genes with low identity to best match ('*') | Matches including missing genes ('-') | Matches including additional genes ('+') | Matches with KL in >1 piece ('?') |
| --- | --- | --- | --- | --- | --- | --- |
| Perfect | 48 (18.2%) | 0 | 0 | 0 | 0 | 0 |
| Very high | 170 (64.4%) | 13 <sup>1</sup> | 25 | 0 | 0 | 0 |
| High | 37 (14%) | 7 <sup>1</sup> | 1 | 37 <sup>2</sup> | 0 | 0 |
| Good | 2 (0.8%) | 0 | 0 | 2 <sup>2</sup> | 1 <sup>3</sup> | 1 |
| Low | 1 (0.4%) | 0 | 0 | 1 <sup>2</sup> | 1 <sup>3</sup> | 0 |
| None | 6 (2.3%) | 0 | 3 | 6 <sup>2</sup> | 4 <sup>3</sup> | 0 |
| Total | 264 | 20 | 29 | 46 | 6 | 1 |

<sup>1</sup> all length discrepancies are confirmed as additional IS insertion(s)
<sup>2</sup> all genes reported 'missing' are due to SNPs predicting frameshifts or due to better
sequence matches of the same gene found a different KL reference sequence in the database
<sup>3</sup> all genes reported 'extra' are better sequence matches of the same gene found a different KL
reference sequence

Seven of the 264 completed genomes were records for the *A. baumannii* reference strain, ATCC 17978 (Table 4), known to carry the KL3 locus [11]. The ATCC 17978 genome was originally sequenced via pyrosequencing [51] and first made available in 2007 under GenBank accession number CP000521 (NCBI assembly number GCA_000015425.1). However, this assembly was later found to include errors [52] and/or gene frameshifts [11], and was subsequently re-sequenced using a combination of Illumina short read and PacBio long read data (GenBank accession number CP012004.1; NCBI assembly number GCA_001077675.1). Here, the re-sequenced genome was assigned to KL3 with ‘perfect’ confidence, whereas the original sequence has a confidence score of ‘high’ with the previously reported KL gene frameshifts resulting in missing genes (‘-’) detected.

**Table 4.** Complete genome sequences for *A. baumannii* strain ATCC 17978.

| Genome assembly<br>accession number | Best<br>match<br>locus | Match<br>confidence | Problems | Sequence<br>coverage | DNA<br>identity | Length<br>discrepancy | Sequencing<br>platform(s) | Assembly and/or<br>polishing software | Read<br>coverage | Upload<br>date |
| --- | --- | --- | --- | --- | --- | --- | --- | --- | --- | --- |
| GCA_000015425 <sup>1</sup> | KL3 | High | -<sup>3</sup> | 100.00% | 99.98% | 0 bp | high-density<br>pyrophosphate<br>DNA<br>sequencing | Pyrosequencing<br>assembly | 21.1X | 2007 |
| GCA_001077675 <sup>2</sup> | KL3 | Perfect |  | 100.00% | 100.00% | 0 bp | Illumina;<br>PacBio | SPAdes v. 2.5.0;<br>HGAP v.<br>2.2.0.133377-patch-3 | 153X | 2015 |
| GCA_001593425 | KL3 | Perfect |  | 100.00% | 100.00% | 0 bp | Illumina MiSeq | Geneious v. 9.1.5 | 300.0x | 2016 |
| GCA_004797155 | KL3 | Perfect |  | 100.00% | 100.00% | 0 bp | PacBio | PacBio SMRT<br>Analysis v. 2.3.0 | 247.19x | 2019 |
| GCA_014672775 | KL3 | High | -<sup>3</sup> | 100.00% | 100.00% | -1 bp | PacBio RSII | HGAP v. 3.0 | 399.24x | 2020 |
| GCA_013372085 | KL3 | Perfect |  | 100.00% | 100.00% | 0 bp | Illumina HiSeq;<br>Oxford<br>Nanopore<br>MiniION | Unicycler v. 0.4.2 | 80.0x | 2020 |
| GCA_011067065 | KL48 | Very high |  | 100.00% | 96.76% | -2 bp | PacBio | Pacbio v. 20K | 231.08x | 2020 |

<sup>1</sup> Genome sequence excluded from RefSeq database due to poor sequence quality
<sup>2</sup> Re-sequenced genome reported in [52]
<sup>3</sup> ‘-‘ is Kaptive problem score indicating missing gene(s)

Three further sequenced versions of the ATCC 17978 genome were assigned a ‘perfect’ match to KL3, whereas another that was sequenced and assembled using only PacBio technology was assigned a ‘high’ KL3 match (Table 4). The remaining assembly (NCBI assembly number GCA_011067065) was a ‘very high’ match to KL48. As this is inconsistent with the previous finding of KL3 in ATCC 17978, the assembly was aligned to the ATCC 17978 reference sequence (GCA_001077675) and found to have only 87% sequence coverage with 98% nucleotide sequence identity. Further inspection using MLST revealed that the genome belongs to sequence type ST2 in the Institut Pasteur scheme, rather than ST437 as for all other ATCC 17978 genome sequences. This suggests that GCA_011067065 is incorrectly named in the GenBank record as ATCC 17978.

### General features of *A. baumannii* K locus sequences

Characterisation of the 237 distinct CPS biosynthesis gene clusters affords the opportunity to re-examine common features of sequences found at the K locus in *A. baumannii* genomes. Amongst the 237 KL, sequence lengths varied between 18.5 kb and 36.8 kb with a mean length of 25 kb (Fig. 4A). The total number of open reading frames (ORFs) per KL also varied, ranging between 16 and 31, with the majority of KL (∼65%) including 20 to 23 ORFs (Fig. 4B). The size of the locus correlated with the number of ORFs present, and the smallest carried no modules for the synthesis of complex sugars. The larger gene clusters generally included larger or more gene modules for complex sugar biosynthesis rather (see Fig. 1 and Fig. 2). For example, large gene modules are required for synthesis of non-2-ulosonic acids such as 5,7-di-*N*-acetylacinetaminic acid that requires 10 genes [53].

**Figure 4.**
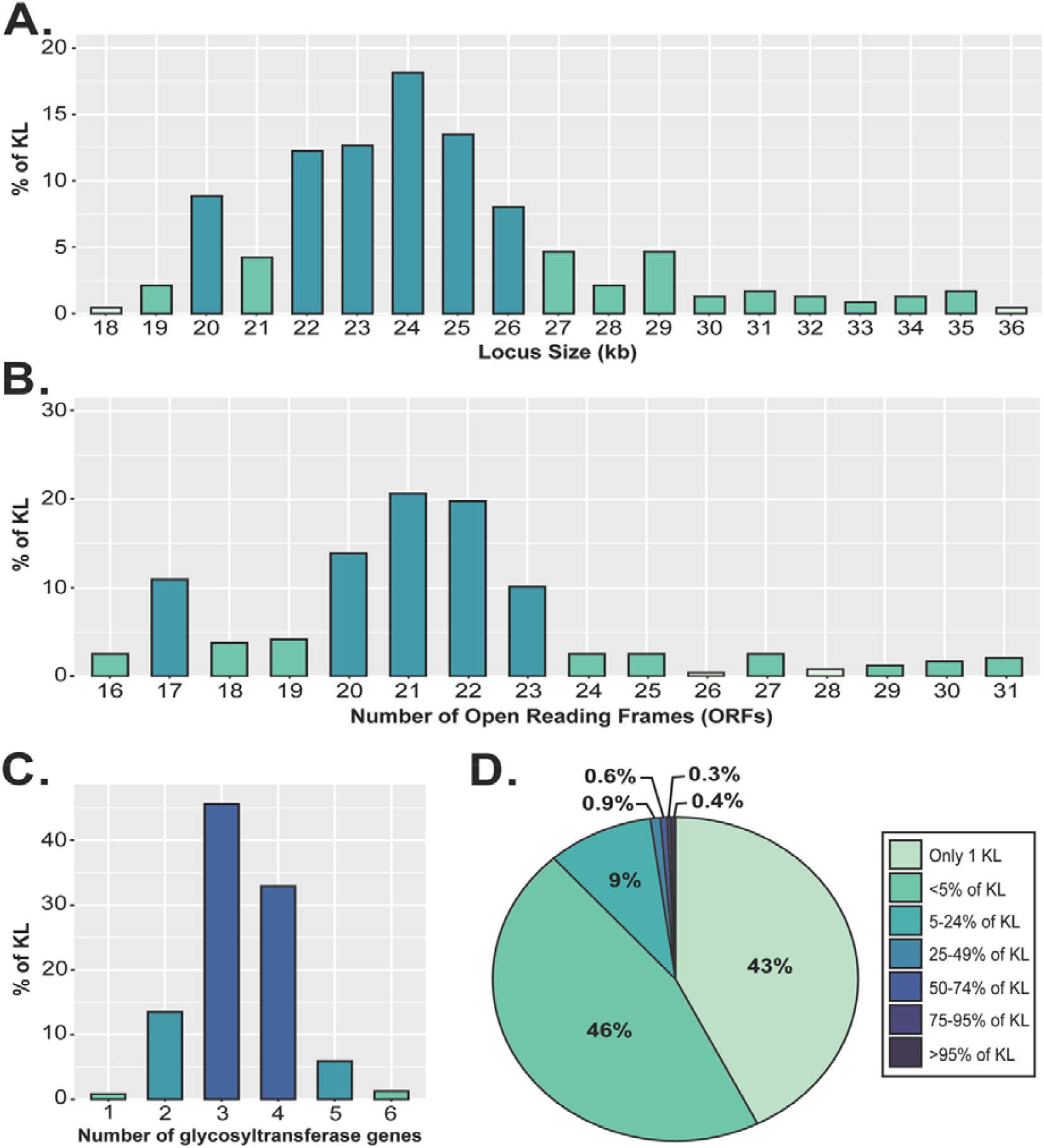
General features of the 237 K loci identified in *A. baumannii* genomes. **(A)** Percentage of 237 KL with specific total sequence length (kb). **(B)** Percentage of 237 KL with specific number of Open Reading Frames (ORFs). **(C)** Percentage of 237 KL with specific number of glycosyltransferase genes (*gtrs*). **(D)** Frequency of 681 gene types (homology groups) found at the K locus across 237 KL. Figures were created in RStudio [46].

A small group of five KL (KL92, KL99, KL142, KL143 and KL145), have a configuration considered unusual for *A. baumannii* as they include a novel segment in Region 2 that includes *itrA4* and *wzi*_*KL*_ genes. Previously, this segment was suggested to have been acquired from a source outside of *A. baumannii* [54].

All KL sequences included between 1 and 6 genes predicted to encode glycosyltransferases, with most KL carrying 3 or 4 *gtr* genes (Fig. 4C) suggesting tetrasaccharide and pentasaccharide K-units are common in *A. baumannii* CPS. The correlation between the number of Gtr encoded and the number of sugars in the K unit has been supported by structural studies with exceptions only for CPS containing L-rhamnose [16, 17, 54].

### Repertoire of genes included in the database

To understand the diversity and distribution of CPS biosynthesis genes across the 237 KL, all genes were grouped into clusters of homologous gene groups using Roary with a cut-off parameter of 85% aa minimum identity for the products, and the groups were used to calculate frequency. This revealed a total of 681 different gene homology groups found across the 237 KL, of which 42.6% of genes were found only in one KL and a further 45.82% found in 2-12 KL (i.e. <5%; Fig. 4D). Nine gene groups occurred in 164 or more gene clusters (>69.2% of KL) and only 1 gene, *pgm*, was found to encode products of >85% aa identity for all 237 KL (100%). This finding was unexpected, as all CPS gene clusters described for *A. baumannii* to date include the same eight genes: *wza, wzb*, and *wzc* genes in ‘Region 1’, *gna* in ‘Region 2, and *galU, ugd, gpi*, and *pgm* genes in ‘Region 3’ [11]. Hence, further assessment of gene product homology groups at the K locus was undertaken.

### Variation in the eight genes always present at the *A. baumannii* K locus

With the exception of *pgm*, 2-4 homology groups were detected for each of the other seven genes that are always present at the K locus, indicating these genes are not completely conserved. The occurrence of >1 homology group for common genes may be due to multiple imports of the same genes into the species via homologous recombination resulting in a change of KL sequence. Sequence diversity in *wza, wzb* and *wzc* genes had been observed previously [12]. However, the level of variation detected here for these genes, as well as for other common genes, is significant and suggests a complex evolutionary history for the *A. baumannii* K locus. For example, two homology groups were found for both the *ugd* and *gpi* genes; one group is present in 97.9% of KL and the smaller group (in 2.1% of KL) occurs in the KL92, KL99, KL142, KL143 and KL145 group described above. However, while 85% aa identity is the cut-off used to define new gene types in *A. baumannii* K loci, variants of these common genes are not currently numbered. Nor are they considered in assignment of new numbers to KL.

For *gna*, three sequence groups had previously been reported in the same position at the beginning of Region 2 [11]. Two of the three types were shown to form a module with either *gne2* (*gna1*) for synthesis of D-GalNAcA or *dgaABC* (*gna3*) for synthesis of D-GlcNAc3NAcA. A third type (*gna2*) is present in all other KL, though its role in CPS production is still unknown. Here, the same three *gna* homology groups were found.

### Further variation in Region 3

When present, the *gne1* gene is located between *gpi* and *pgm*. This gene is often present and is required for synthesis of UDP-D-GalNAc and/or UDP-D-Gal [11]. Though the presence of a small variable portion of Region 3 between the *gpi* and *pgm* genes has been previously reported [11, 12], diversity in this region had not been further investigated. Here, a total of 217 of 237 KL (91.56%) were found to carry additional coding sequence(s) between *gpi* and *pgm*. A *gne1* gene was present on its own in 95 KL (40.08%), though a further 97 KL (40.93%) included *gne1* adjacent to additional *pgt1, pgt2, pet1* or *atr* genes (Table 5). A further 23 KL include only *pgt1* between *gpi* and *pgm*. Besides *gne1*, a role for all other genes found between *gpi* and *pgm* in CPS biosynthesis has not yet been established.

**Table 5.** Number of KL with different gene combinations between *gpi* and *pgm* in Region.

| **Gene combination** | **Number of KL** |
| --- | --- |
| *gne1* only | 95 |
| *gne1/pgt1* | 76 |
| *gne1/pgt2* | 1 |
| *pgt1* only | 23 |
| *gne1/pet1/orf/orf* | 3 |
| *gne1/orf/atr32/atr33* | 3 |
| *gne1/atr15* | 2 |
| *gne1/orf/atr20* | 1 |
| *gne1/atr42/atr43* | 6 |
| *gne1/atr24* | 1 |
| *gne1/orf/atr5-like* | 1 |
| *gne1/atr12* | 3 |
| None | 22 |
| **TOTAL** | **237** |

As the variation at this position in Region 3 is known to affect CPS structure only if D-Gal*p* or D-Gal*p*NAc are present, some groups of KL are likely to produce the same CPS structure. This has been the case for KL2/KL81 and KL3/KL22 pairs that are known to produce the same structure [23]. The gene clusters in these two pairs differ from each other only in the presence of a *pgt1* gene between *gpi* and *pgm*, which has no defined role in CPS biosynthesis. A further 21 examples of pairs or groups of KL that differ in the presence/absence of *pgt1* and/or other genes found between *gpi* and *pgm* in Region 3 were detected amongst the 237 KL (listed in Table 6). Comparisons of some of these pairs or groups can be seen in published KL compilations [15, 18, 25]. Hence, it is possible that further examples of KL that produce the same K type may be found as more CPS structures are determined. Of these 21, 15 groups include one KL with associated structural data (bold in Table 6), possibly representing the structure produced by the other KL of the same group.

**Table 6.**
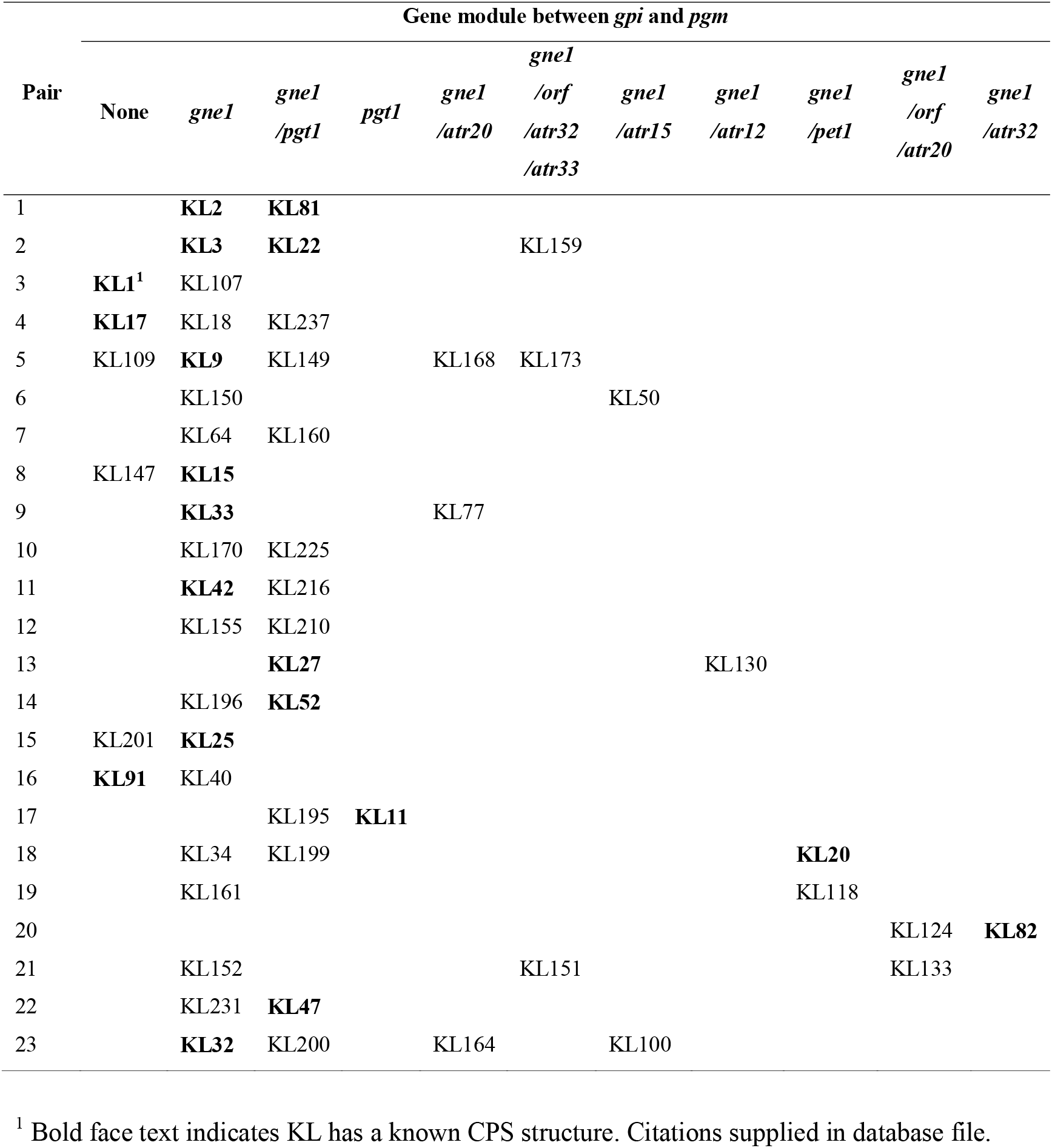
K loci predicted to produce the same CPS structure.

| Pair | Gene module between *gpi* and *pgm* |  |  |  |  |  |  |  |  |  |  |
| --- | --- | --- | --- | --- | --- | --- | --- | --- | --- | --- | --- |
|  | None | *gne1* | *gne1*<br>*/pgt1* | *pgt1* | *gne1*<br>*/atr20* | *gne1*<br>*/orf*<br>*/atr32*<br>*/atr33* | *gne1*<br>*/atr15* | *gne1*<br>*/atr12* | *gne1*<br>*/pet1* | *gne1*<br>*/orf*<br>*/atr20* | *gne1*<br>*/atr32* |
| 1 |  | **KL2** | **KL81** |  |  |  |  |  |  |  |  |
| 2 |  | **KL3** | **KL22** |  |  | KL159 |  |  |  |  |  |
| 3 | **KL1**<sup>1</sup> | KL107 |  |  |  |  |  |  |  |  |  |
| 4 | **KL17** | KL18 | KL237 |  |  |  |  |  |  |  |  |
| 5 | KL109 | **KL9** | KL149 |  | KL168 | KL173 |  |  |  |  |  |
| 6 |  | KL150 |  |  |  |  | KL50 |  |  |  |  |
| 7 |  | KL64 | KL160 |  |  |  |  |  |  |  |  |
| 8 | KL147 | **KL15** |  |  |  |  |  |  |  |  |  |
| 9 |  | **KL33** |  |  | KL77 |  |  |  |  |  |  |
| 10 |  | KL170 | KL225 |  |  |  |  |  |  |  |  |
| 11 |  | **KL42** | KL216 |  |  |  |  |  |  |  |  |
| 12 |  | KL155 | KL210 |  |  |  |  |  |  |  |  |
| 13 |  |  | **KL27** |  |  |  |  | KL130 |  |  |  |
| 14 |  | KL196 | **KL52** |  |  |  |  |  |  |  |  |
| 15 | KL201 | **KL25** |  |  |  |  |  |  |  |  |  |
| 16 | **KL91** | KL40 |  |  |  |  |  |  |  |  |  |
| 17 |  |  | KL195 | **KL11** |  |  |  |  |  |  |  |
| 18 |  | KL34 | KL199 |  |  |  |  |  | **KL20** |  |  |
| 19 |  | KL161 |  |  |  |  |  |  | KL118 |  |  |
| 20 |  |  |  |  |  |  |  |  |  | KL124 | **KL82** |
| 21 |  | KL152 |  |  |  | KL151 |  |  |  | KL133 |  |
| 22 |  | KL231 | **KL47** |  |  |  |  |  |  |  |  |
| 23 |  | **KL32** | KL200 |  | KL164 |  | KL100 |  |  |  |  |

<sup>1</sup> Bold face text indicates KL has a known CPS structure. Citations supplied in database file.

### Genes for biosynthesis of sugars and addition of substituents

The number of homology groups relating to specific modules of genes for sugar biosynthesis or functional categories of gene products were manually curated to gain insights into the possible diversity in sugars and non-sugar substituents that can be incorporated into *A. baumannii* CPS. A total of 34 possible modules of gene(s) for the synthesis of complex sugars (described in Table 7) were found. Of these modules, 29 have been reported previously and three are variants of known modules. Two modules, *rhnABC* and *gnlAB*, are described here for the first time (see above). While most KL include at least one gene module for complex sugar biosynthesis, 30 KL did not include any sugar synthesis gene module(s) in Region 2 and these are likely to produce CPS with neutral sugars [23, 26, 29, 37] synthesised by common genes in Region 3 [11]. The *rmlBDAC* module for L-Rhamnose synthesis is the most common sugar gene module across the 237 KL types (found in 15.6%), though gene modules for synthesis of sugars belonging to the non-2-ulosonic acid family (i.e. *psa, lga, aci, ela*, and *neu*) were collectively found in 29.11% of KL.

**Table 7.**
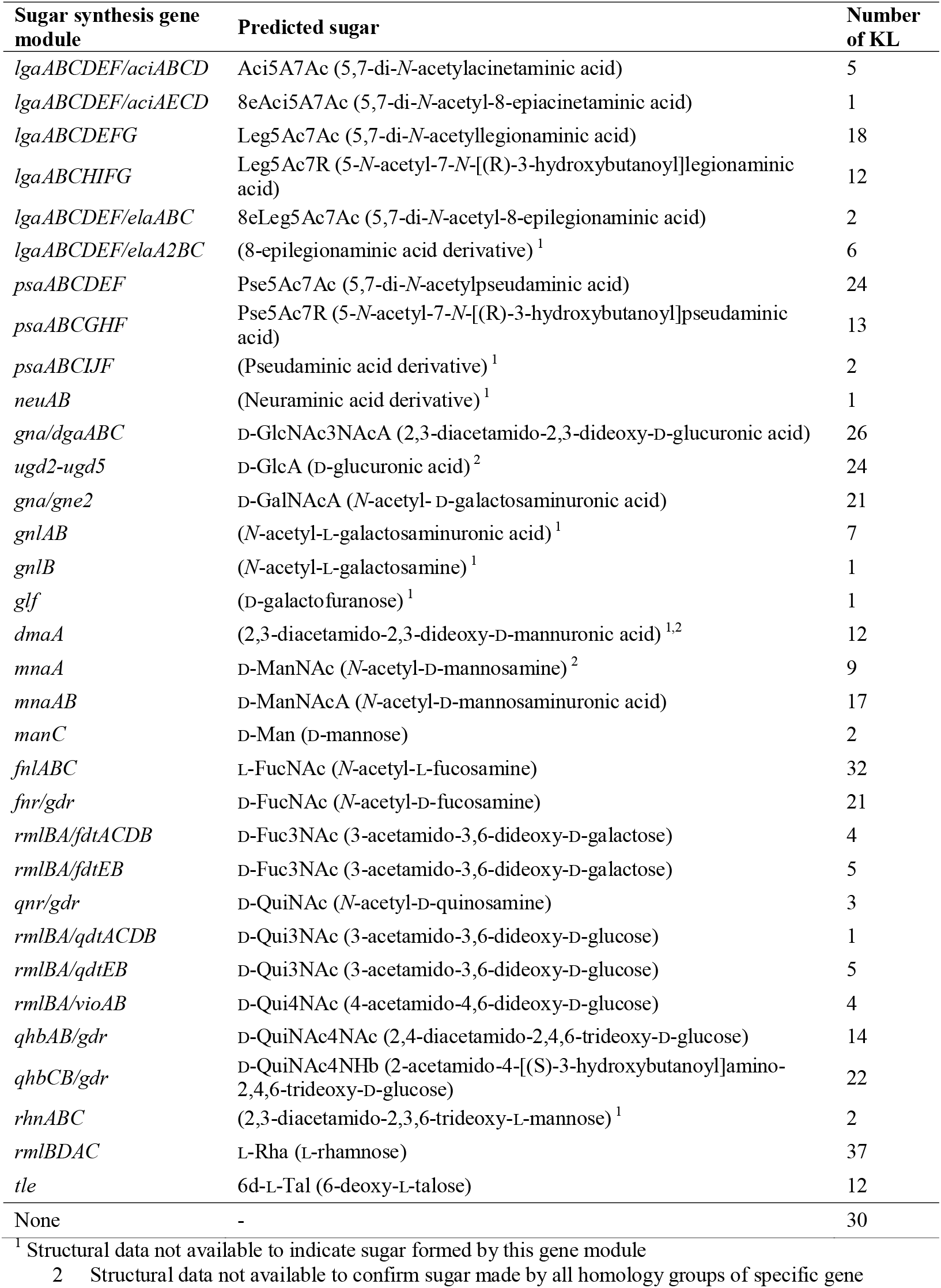
Different sugar synthesis gene modules identified in Region 2 across 237 KL.

In addition to complex sugar biosynthesis genes/modules, 40 different genes for modifying a monosaccharide in the K unit via the addition of a substituent were identified in Region 2. These included 31 *atr* genes for the transfer of acyl-/acetyl groups, 8 *ptr* genes for the addition of pyruvate, and 1 *alt* gene for transfer of L-alanine. The presence of 34 sugar biosynthesis gene modules and 40 different genes for the addition of non-sugar substituents to the CPS therefore indicates significant potential for diversity in sugar composition and K-unit decoration of *A. baumannii* CPS structures.

### Genes for initiating transferases and glycosyltransferases

Biosynthesis of *A. baumannii* CPS in the cytoplasm is known to begin with the transfer of a ‘first’ sugar to a lipid carrier in the inner membrane, and six possible first sugars transferred by one of six distinct initiating transferases (Itrs) belonging to one of two families (ItrA or ItrB) are known [55]. A seventh non-functional Itr type, ItrB2, has also been reported [11]. Here, the same seven *itr* genes (*itrA1-itrA4* and *itrB1-itrB3*) were found across the 237 KL as expected with no new types identified. The *itrA2* and *itrA3* genes were found to be the most common, present in 91 and 81 different KL, respectively. This indicates that UDP-D-GalNAc (ItrA2) and UDP-D-GlcNAc (ItrA3) are common first sugar substrates for *A. baumannii* CPSs, consistent with what has been observed with available structural data.

Following the addition of the first sugar to the lipid carrier, CPS biosynthesis progresses with the addition of further monosaccharides to the first sugar to build a complete K-unit oligosaccharide [11]. The glycosidic linkages between sugars are formed by glycosyltransferase enzymes that are encoded by either *gtr* or *kpsS* genes at the K locus. As the specificity of Gtr/KpsS enzymes for their sugar donor and acceptor substrates can vary, numbering new types is important. While new numbers are currently assigned to genes using a cut-off of 85% aa identity, evidence has emerged that some Gtrs that share <85% aa identity can form the same linkage as one another [17], while others sharing >85% aa identity can form different linkages [48, 56]. Nonetheless, differentiation between different Gtr/KpsS types can provide insights into diversity in the linkages possible between sugars in the K-unit. Using the cut-off of 85% aa identity, a total of 272 homology groups (267 *gtr* and 5 *kpsS*) were found across the 237 KL. Further work will be needed to analyse Gtr sequences to identify relationships between them and how these relate to linkages formed.

### Genes for K-unit and CPS processing

Region 2 usually also harbours *wzx* and *wzy* genes required for processing oligosaccharide units to form long chain polymers as part of the Wzy-dependent pathway for CPS biosynthesis [12]. Wzx is the translocase that flips oligosaccharide units into the periplasm for polymerisation into chains by the Wzy polymerase. Across the 237 KL, a total of 81 *wzx* and 137 *wzy* gene groups were found. It is unclear what effect *wzx* sequence diversity has on CPS biosynthesis, though the variety of *wzy* genes may reflect many different linkages possible between oligosaccharide units in the CPS polymer. With the exception of the *wzy*_KL8_ type (Fig. 1B) which is located to the left of the *wza-wzb-wzc* genes, and the three (KL19, KL24 and KL39) that do not contain any *wzy*, all *wzy* groups were found in Region 2. The absence of *wzy* in these three gene clusters has been described previously, where Wzy function is supplemented by a *wzy* gene located in genomic islands present elsewhere in the chromosome [35, 36]. However, a relative of KL24, KL146, that does include a *wzy* gene was recently identified [4] indicating that the original *wzy* gene may have been lost. Interestingly, two KL (KL67 and KL134) include 2 distinct *wzy* genes, though it is not known if both contribute to CPS assembly. As *wzx* and *wzy* genes can be shared by K loci, to distinguish *wzx* and *wzy* groups for future typing, a suffix was added to each *wzx* and *wzy* type (defined by the 85% aa identity cutoff) indicating the name of the first KL a group was identified in.

Finally, CPS assembly on the cell surface is mediated by a Wzi outer membrane protein encoded by a *wzi* gene located outside the K locus [55]. However, in rare cases, a second *wzi* type has been found at the K locus [54], and is referred to as *wzi*_*KL*_. This *wzi* gene is co-located with *itrA4* and was found only in the 5 KL that carry different *ugd* and *gpi* genes (see above).

### Abundance of KL and K types in the genome pool

To examine the ability of the updated database to detect diversity, the frequency of KL and K types detected across the 8994 genome assemblies was calculated. An assessment of best match loci reported by *Kaptive* with a confidence score of ‘good’ or above, regardless of detected ‘problems’ or possible discontiguous locus sequence, revealed that 19 KL were not found amongst the 8994 genome assemblies studied. Two of these were KL38 and KL78 that are no longer considered *A. baumannii* sequences, and 17 KL that are currently only available as short reads or extracted locus sequences in the GenBank non-redundant database. Some KL were overrepresented, with 17 KL each found in >1% of genome assemblies (Fig. 5A) and collectively representing 74.6% of the total genome pool. The other 25.4% of genome assemblies included 201 KL. KL2 was found to be the most common K locus sequence found in 16.5% of genomes, while KL3 and KL22, both of which produce a K3 type CPS, together represent 20% of genomes.

**Figure 5.**
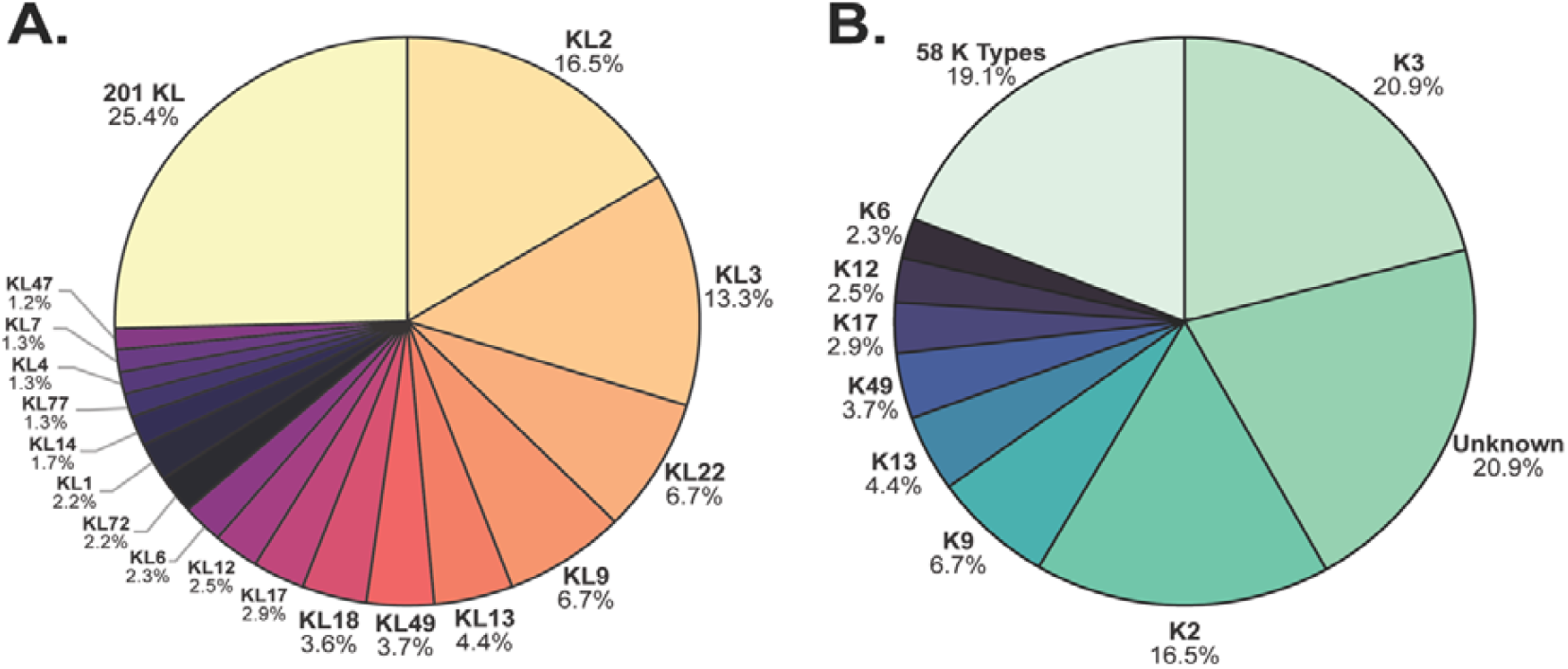
Distribution of **(A)** best match locus and **(B)** best match type amongst the 8994 genome assemblies assigned a confidence score of ‘good’ or above by *Kaptive*. Figures were created in RStudio [46].

A similar assessment of K type demonstrated that 66 K types could be predicted from the assemblies reported with a confidence score of ‘good’ or above. Eight K types were represented in >1% of genomes each (Fig. 5B), whereas 58 K types represented 19.1% and the remaining 20.9% of matches were listed as ‘unknown’ because relevant structures are not available. As the KL3 and KL22 gene clusters are known to produce the same K3 type structure [23], K3 was found to be the most common consistent with the high proportion of KL3 and KL22 in the genome pool followed by K2 (Fig. 5A).

## 8. Discussion

In this study, we provide a comprehensive update to the *A. baumannii* CPS gene reference sequence database, providing 237 fully annotated K loci sequences and six ‘extra’ gene loci outside of the K locus. Each KL record has linked structural information to enable ‘type’ to be inferred wherever possible based on detection of specific KL with or without ‘extra’ genes based on special logic parameters. However, since the conclusion of this work, an additional 4 novel KL have been identified in recently reported *A. baumannii* genomes. Hence, a total of 241 distinct KL sequences have been released into the *Kaptive v. 2*.*0*.*1 A. baumannii* CPS gene reference sequence database. As we did not further investigate genomes with K loci detected in more than 1 piece in this study, we expect there may be more sequences yet to be documented amongst the 8994 genome assemblies included in this study.

Of the 6726 genome assemblies with a KL match found in a single piece, 98.22% were able to be assigned matches with either ‘perfect’, ‘very high’ or ‘high’ confidence demonstrating the utility of the database to type KL in *A. baumannii* genomes. However, for the majority of ‘high’ matches, *Kaptive* had detected missing gene products that appeared to be the result of frameshifts in gene sequence(s). Further work will be needed to determine if these frameshifts are the result of sequence/assembly errors, or whether frameshifts are real and if they give rise to changes in the CPS composition therefore warranting a new KL designation to distinguish mutants from wildtype sequences. Given that our inspection of 264 complete genome sequences suggests that the quality of both the sequence and the assembly influences the confidence score, the remaining genome assemblies that were not assigned matches with either ‘perfect’, ‘very high’ or ‘high’ confidence, may be of poorer quality. Nonetheless, KL variants were found in the genomes with confidence scores of ‘very high’ or lower, and novel KL were found amongst assemblies with ‘high’, ‘good’, ‘low’ or even ‘none’ confidence levels. Hence, users are encouraged to check sequence/assembly quality prior to KL typing, and manually inspect any assembly assigned a *Kaptive* match less than ‘perfect’.

With the addition of ‘type’ to the update, users are now also able to infer structural properties of the CPS from their genome data when this is available or can be reasonably inferred. Whilst the inclusion of >237 KL significantly enhances the ability to predict CPS type using WGS, structural studies are currently the only definitive way to determine if genes outside the K locus contribute to the determination of a specific CPS structure. Hence, further examples of genes elsewhere in the genome are likely to be found as more structural data coupled with KL gene content analysis becomes available. Though further work is needed to improve the predictive power of WGS for determination of CPS type, the database is a valuable tool for epidemiological studies. Our analysis showed that 17 KL represent a large proportion of the genome assemblies included in this study. Though this may be due to bias in the dataset associated with strain sampling for outbreak studies, particularly the over-representation of genome for GC2 isolates, new KL described here were each found in <1% of genomes in the pool, indicating that the previous iteration of the database had captured the most common KL.

Additional analysis of K locus gene repertoire revealed a total of 681 gene types amongst 237 KL, including 601 (88.3%) genes found in <5% of KL and 286 of these (42%) present in only one KL. Interestingly, 95 of 237 KL include one or more unique genes, suggesting that new gene clusters may arise by acquisition of novel genes, as well as by the reassortment or exchange of genes between already established KL. With the K locus known to undergo recombination [57], sequences of shared genes are likely to exhibit a degree of sequence variation. However, the finding that 7 of the 8 genes always present at the K locus included 2 or more product homology groups of <85% aa identity was unexpected, suggesting that multiple imports of these genes into the species has occurred over time. While new KL numbers are usually assigned to any new combination of genes at the K locus defined by an 85% aa identity cut-off in one or more gene products, new numbers are not warranted for KL that differ from a reference sequence only in one or more of these 8 genes. Hence, we continue to assign new numbers only for any new combination of genes in Region 2 (between *gna* and *galU*) and the variable portion of Region 3 (between *gpi* and *pgm*), regardless of whether a difference in CPS structure is expected. As structural data for corresponding CPS continues to grow, examples of KL that produce the same CPS structure will likely be uncovered.

Gene repertoire analysis further identified genes for 34 different complex sugars, >400 for glycosidic linkages (*gtr, kpsS* and *wzy*) and >40 for K-unit modifications (*atr, alt*, and *ptr*) across 237 KL sequences, predicting substantial variation in CPS sugars, sugar linkages, and also acetyl-, acyl-, pyruvyl, and L-alanine decorations. This extraordinary diversity observed in CPS biosynthesis genes complicates next-generation therapies, including vaccines and bacteriophage strategies. However, fine-scale analysis of individual CPS structures and their specific biosynthesis genes can inform tailored approaches to alternate patient treatments. The increase in the number of KL and inclusion of K type information in the updated reference database will significantly enhance epidemiological tracking efforts and assist with building a comprehensive understanding of the circulation of important strains both locally and globally.

## 9. Author statements

### 9.1 Authors and contributors

Conceptualization, JJK; Data curation, JJK; Formal analysis, SMC and JJK; Funding acquisition SMC, RMH and JJK; Investigation, SMC, RMH and JJK; Methodology, SMC and JJK; Visualization, SMC and JJK; Supervision, RMH and JJK; Writing – original draft, SMC and JJK; Writing – review & editing, RMH and JJK

### 9.4 Ethical approval

N/A

### 9.5 Consent for publication

N/A

## 9.6 Acknowledgements

We thank Kelly Wyres and Ryan Wick from Monash University, Australia, and Kathryn Holt from the London School of Hygiene & Tropical Medicine, UK, for their assistance with updating the database on the *Kaptive* platforms.

